# Live soil ameliorated the negative effects of biodegradable but not non-biodegradable microplastics on the growth of plant communities

**DOI:** 10.1101/2023.10.12.562149

**Authors:** Yanmei Fu, Ayub M. O. Oduor, Ming Jiang, Yanjie Liu

## Abstract

1. Plastic pollution has become a global environmental problem. Alternative use of biodegradable plastics has been proposed to mitigate the pollution problem caused by the traditional non-biodegradable plastics but the relative impacts of both types of microplastics on plant community productivity and diversity remain unknown. Microplastics can affect growth of individual plants directly by altering plant physiological processes and indirectly by altering soil biota that in turn influence plant growth. However, it remains unknown whether soil biota can mediate impact of biodegradable and non-biodegradable microplastics on plant community productivity and diversity due to a lack of studies on the topic.
2. Here, we performed a greenhouse experiment with six plant communities and five biodegradable and five non-biodegradable microplastics to test whether: 1) biodegradable microplastics have a less negative effect on plant community biomass production and diversity than non-biodegradable microplastics, and 2) soil microorganisms differentially mediate the effects of non-biodegradable and biodegradable microplastics on plant community biomass production and diversity. We employed a fully crossed factorial design to grow the six plant communities in the presence *vs*. absence of the 10 microplastics individually and in live soil *vs*. sterilized soil.
3. Results show that live soil ameliorated the negative effects of biodegradable microplastics on shoot biomass of the plant communities, but microplastics suppressed plant community diversity more strongly in live soil than in sterilized soil regardless of microplastics types under averaged across all treatments. Furthermore, the specific microplastics polymers were the main drivers of these results.
4. *Synthesis and applications:* Overall, our findings indicate that even biodegradable microplastics, e.g. PBS, which are considered environmentally friendly, still pose significant ecological risks to the structure and productivity of plant communities with potential implications for functioning of terrestrial ecosystems. Future studies may identify the specific taxa of soil microorganisms that may have degraded the microplastics that we studied, their rates of biodegradation, and the effects thereof on plant community structure and productivity under more natural field conditions in contrasting climatic conditions.

## Introduction

Plastic pollution has become a global environmental problem (Rezania *et al*. 2018; Wang *et al*. 2020). Although most countries have issued restrictions on the use of plastics to control plastic pollution (Raha, Kumar & Sarkar 2021), recent research indicates that the growth of plastic waste is expected to exceed efforts to reduce plastic pollution (Borrelle *et al*. 2020). The larger plastics gradually degrade into mesoplastics (5-25 mm) and microplastics (<5 mm) (Andrady 2011; Jabeen *et al*. 2017). Several studies have shown that microplastics occur widely in aquatic ecosystems where they are already affecting aquatic life adversely (Sarker *et al*. 2020; Savoca, McInturf & Hazen 2021; Xu & Yu 2021). However, the impacts of microplastics on terrestrial ecosystems remain comparatively little studied despite evidence suggesting that terrestrial ecosystems are a major sink of microplastics (Xu *et al*. 2020; Baho, Bundschuh & Futter 2021).

Microplastics can enter terrestrial soils from a multitude of sources. For instance, low-density polyethylene and polypropylene, which are used for mulching, are considered key sources of microplastics in agricultural soils (Bläsing & Amelung 2018). Other major sources of microplastics in soils include organic wastes such as compost and sewage sludge containing microplastics that are widely used as soil amendments, wastewater, effluents from wastewater treatment plants and urban runoffs, contaminated river and lake water, littering and illegal dumping of plastic wastes, particles from tire abrasion on roads, abrasion of synthetic textiles during washing, fertilizer and seed coatings, and atmospheric fallout from surfaces such as managed landfills or streets (Ren *et al*. 2018; Yang *et al*. 2021). Generally, degradation of petroleum-based microplastics in the soils is an extremely slow process, which may last hundreds and even thousands of years for full mineralization to occur (Zubris & Richards 2005; Zhou *et al*. 2021). Consequently, the concentration of microplastics in the top layer of some highly polluted soils was found to be as high as 7% of the total soil weight (Fuller & Gautam 2016). Therefore, studies are needed that assess the impacts of plastic pollution on plant growth in terrestrial ecosystems.

To date, studies that have tested the effects of microplastics on plant growth performance have focused on single species, with mixed results. For instance, *Triticum aestivum* (wheat), *Lolium perenne* (perennial ryegrass) (Boots, Russell & Green 2019), *Arabidopsis thaliana* (Sun *et al*. 2020) and *Bacopa sp*. (Yu *et al*. 2021) produced less biomass when grown in media with microplastics than without microplastics, while *Allium fistulosum* (spring onion) exhibited variable growth responses (increases and decreases) to different types of microplastics (de Souza Machado *et al*. 2019). In another study, microplastics inhibited root growth in *Lactuca sativa* (Italian lettuce) and *Zea mays* (corn) but not in *Raphanus sativus* (radish) and *T. aestivum* (Gong *et al*. 2021). The variable responses of test plants observed in the single-species studies suggest that within plant communities, different plant species may be affected to different degrees by microplastics, which could potentially affect community productivity and structure (Lozano & Rillig 2020). In a plant community, some species may take advantage of soil conditions altered by microplastics while other species may have depressed growth (Lozano & Rillig 2020), which may cause community diversity loss and compositional shifts. However, studies are lacking that test how microplastics may affect plant community productivity and diversity. To the best of our knowledge, only two studies have tested the effects of microplastics on plant community performance, and they focused only one type of microplastic (Lozano & Rillig 2020; Deng *et al*. 2022). A prediction of impacts of microplastic pollution on plants requires studies that employ a wide range of microplastic types under more natural plant community settings.

Microplastics can affect plant growth directly by impacting plant physiology and indirectly through alteration of soil physical, chemical, and biological properties that then feedback to influence plant growth (Baho, Bundschuh & Futter 2021). For example, an experiment demonstrated that *A. thaliana* seedlings directly absorbed and accumulated nanoscale (<100 nm) microplastics, which then directly altered physiological processes and growth performance of the seedlings (Sun *et al*. 2020). Separately, microplastics sized 1 µm were shown to enter roots of *Daucus carota* (carrot) and accumulate in the intercellular spaces, which caused the carrot to lose crispness (Dong *et al*. 2021). The absorbed microplastics can induce general oxidative stress responses, which influence expression (upregulation and downregulation) of certain genes such as those that encode antioxidant enzymes and regulate production of defence compounds and growth (Jiang *et al*. 2019; Sun *et al*. 2020). Microplastics can alter soil properties, including, distribution of water-stable aggregates, bulk density, water-holding capacity, pH, and inorganic mineral content as well as activities and diversities of soil biota (Liu *et al*. 2017; de Souza Machado *et al*. 2018; Boots, Russell & Green 2019; Liang *et al*. 2019; Zang *et al*. 2020). Such changes in soil properties may impact nutrient and water absorption by plants. For example, adsorption of microplastics on root hairs may reduce uptake and transport of water and nutrients by plants resulting in a decrease in plant growth (Sun *et al*. 2020). Plant growth performance depends on activity of soil microorganisms that act as symbionts (e.g., nitrogen-fixers and mycorrhizal fungi) and antagonists such as pathogens (Kuzyakov & Xu 2013; Wagg *et al*. 2014; de Souza Machado *et al*. 2018). Therefore, as activities and diversities of soil microorganisms are sensitive to changes in soil physical and chemical properties (Karimi *et al*. 2020), microplastics can affect plant growth indirectly by altering soil microbial habitats (Zubris & Richards 2005; Rillig *et al*. 2019; Li *et al*. 2021). Microplastics and their additives (e.g., phthalate acid esters) may also have direct chemical toxicity on soil microorganisms (Wang *et al*. 2016). Empirical evidence shows that microplastics can affect microbe-driven soil nitrogen cycling and enzyme activity of soil microbes (Chen *et al*. 2020; Yi *et al*. 2021). Therefore, studies are needed that investigate how microplastics affect plant community productivity and diversity indirectly through soil microbial activities.

In the recent past, alternative use of biodegradable plastics has been proposed to mitigate the pollution problem caused by the traditional non-biodegradable plastics (Rahman & Bhoi 2021). However, information on the potential risks associated with biodegradable microplastics in the terrestrial ecosystems is still scarce (Zuo *et al*. 2019). Thus, it remains questionable whether biodegradable plastics are as much of a “guarantee” against the impact of plastic pollution on terrestrial ecosystems as one might expect (Sintim & Flury 2017). There are no empirical results showing that biodegradable microplastics can be completely degraded in a fairly short period of time (e.g., one crop growth season per year) (Agarwal 2020). Moreover, some studies indicate that biodegradable microplastics with high surface area could concentrate more toxic chemicals (Wang *et al*. 2016; Qi *et al*. 2020), which suggests that biodegradable microplastics may pose a significant risk to terrestrial ecosystems compared to non-biodegradable microplastics. In fact, some studies have shown that biodegradable microplastics do not have a less negative effect than non-biodegradable microplastics. For example, biodegradable plastic residues (both macro-and microplastics) had stronger negative effects on *T*. *aestivum* than non-biodegradable (i.e., polyethylene) plastics (Qi *et al*. 2020). Separately, biodegradable microplastics butylene adipate-co-terephthalate (PBAT) and Polylactic acid (PLA) had stronger negative effects on the growth of *Phaseolus vulgaris* (common bean) than a non-biodegradable microplastic low-density polyethylene (LDPE) (Meng *et al*. 2021). Nevertheless, no study has assessed the relative impacts of biodegradable and non-biodegradable microplastics on plant community productivity and diversity.

The objective of the present study was to test whether biodegradable and non-biodegradable microplastics affect plant community productivity and diversity directly and indirectly through soil microorganisms. Specifically, we performed a greenhouse experiment with six communities of plant species and 10 microplastics (five biodegradable and five non-biodegradable) to address two questions: 1) Do biodegradable microplastics have a less negative effect on plant community biomass production and diversity than non-biodegradable microplastics? 2) Do soil microorganisms mediate the effects of non-biodegradable and biodegradable microplastics on community biomass production and diversity?

## Material and Methods

### Plant species selection

We obtained seeds of 28 randomly selected native plant species from 25 genera and 12 families (**Table S1**). All the species are herbaceous, and commonly co-occur in many grasslands of China. From the pool of 28 species, we constructed six communities of five species each, and each community contained two grass and three forb species (**Table S1**). The communities represent those that occur naturally in grasslands in China. Each species was used not more than twice to create the six communities. The seeds of all species were collected from natural populations in grasslands of China (**Table S1**).

### Microplastic selection

We selected 10 commonly used different types of microplastics, including five biodegradable microplastics and five non-biodegradable microplastics. As biodegradable microplastics, we used Polybutylene succinate (PBS) (Mulvane, Kansas), Polycaprolactone (PCL) (Malmö, Sweden), Polyhydroxyl alkanoates (PHA) (Shenzhen, China), Poly-3-hydroxybutyrate (PHB) (Rialto, California), and Polylactic acid (PLA) (Willich, Germany). As non-biodegradable microplastics, we used Ethylene-vinyl acetate (EVA) (Seoul, South Korea), Poly amide 66 (PA66) (Wilmington, Delaware, USA), Polyethylene-terephthalat (PET) (Barcelona, Spain), Poly oxymethylene (POM) (Wilmington, Delaware, USA), and Polyvinyl chloride (PVC) (Leominster, Massachusetts, USA). All the microplastic polymers were acquired as powder with two different sizes by sieving through 150 µm (PCL, PHA, PHB, PLA, PVC, POM, PA66) and 180 µm (PBS, EVA, PET) meshes. We purchased the microplastics from Suzhou XinSuYu Company in Suzhou, China.

### Experimental set-up

To test soil microbe-mediated effects of biodegradable and non-biodegradable microplastics on plant community biomass production and diversity, we conducted a pot experiment in a greenhouse of the Northeast Institute of Geography and Agroecology, Chinese Academy of Sciences (43°59’49” N, 125°24’03” E). We employed a fully crossed factorial design to grow the six plant communities in the presence *vs*. absence of microplastics and in the presence *vs*. absence of live soil (**Figure 1**). We replicated each treatment combination four times, which resulted in 528 experimental pots: 6 plant communities × 2 soil treatments (live soil *vs.* sterilized soil) × 11 microplastic treatments (five biodegradable microplastics, five non-biodegradable microplastics, no-microplastic) × 4 replicates.

**Figure 1.**
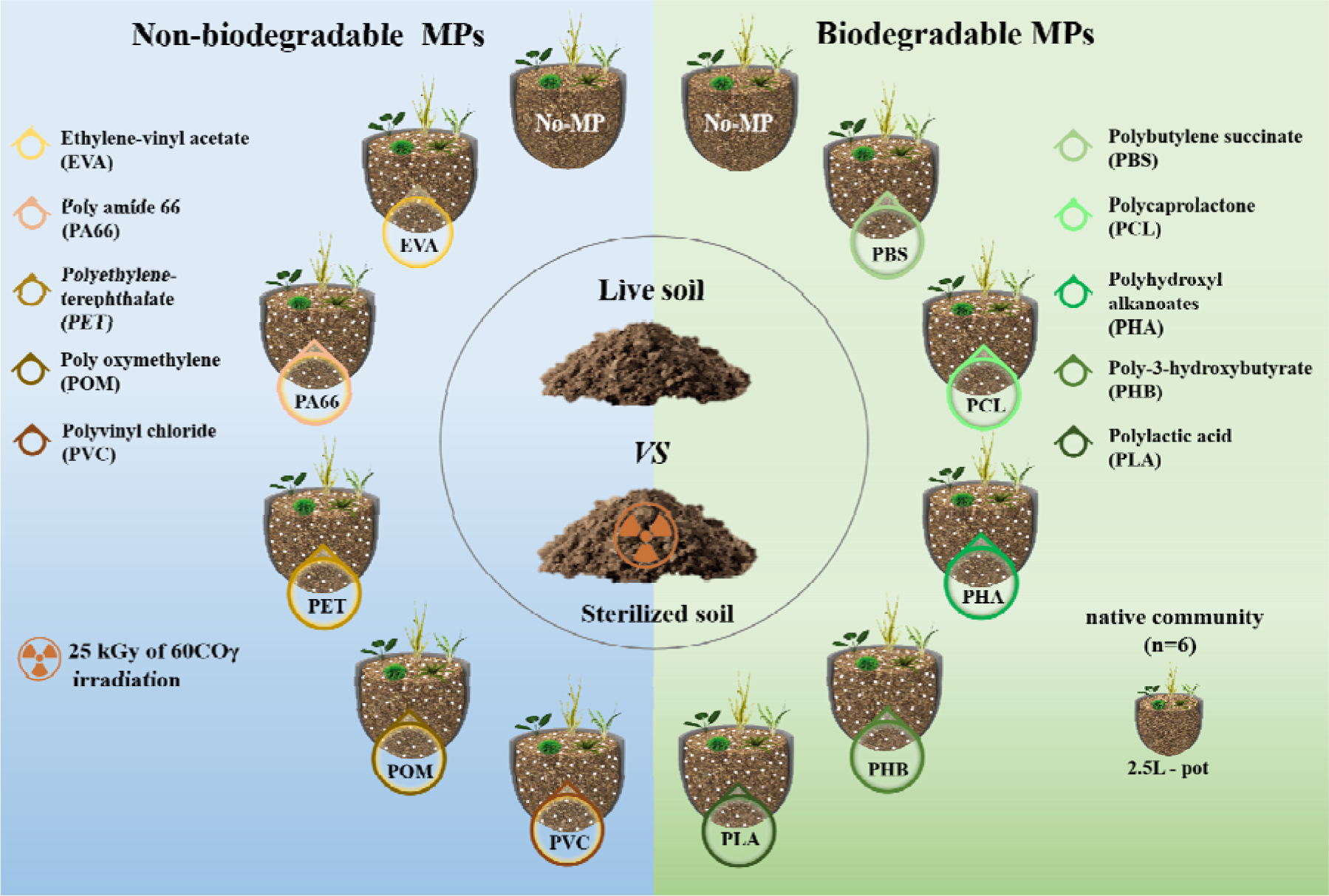
A schematic of the experimental design. No-MP denotes absence of a microplastic (MPs), while differently colored circles represent the addition of different types of microplastics. Each plant community was grown in separate soils that contained the 10 different types of microplastics and in a soil with no microplastic (No-MP). A half of the soil was sterilized to kill soil microorganisms; the live soil represented a live soil inoculum. Each of the six native plant communities was constructed with five species (*cf.* Table S1).

To raise seedlings of the 28 plant species that were used to construct native plant communities, we sowed the seeds of each species separately into trays (12 cm × 12 cm × 4.5 cm) that had been filled with a pre-sterilized (autoclaved at 121°C for two hours) commercial potting soil (Pindstrup Plus, Pindstrup Mosebrug A/S, Denmark) between 7 to 14 August 2020. Prior to sowing, the seeds were surface-sterilized with 75% absolute alcohol for 1 minute, followed by 5-minute soak in 0.5% sodium hypochlorite, and finally rinsed thoroughly with sterile water . As the 28 species have different germination speeds, we sowed the seeds on different dates to ensure that all species were in a similar developmental stage at the start of the experiment (**Table S1**). On 30 August 2020, we used the resultant seedlings to construct six communities of five species each within individual 2.5 L circular plastic pots (top diameter: 18.5 cm, bottom diameter: 12.5 cm, height: 15 cm, 5 holes at the bottom). We transplanted five similar-sized seedlings (each seedling represented one species) from a pool of 28 species above into the pots. The five seedlings were transplanted in a circular formation and spaced at equal distances from each other within a pot. Prior to transplant, each pot was filled with a substrate that comprised 37.5% (v/v) sand, 37.5% (v/v) fine vermiculite, and 25% (v/v) of live or sterilized field soil inoculum. The field soil inoculum was obtained from a grassland site (top 20 cm) (44°35′40.78″N, 123°30′50.94″E) on 21 August 2020. Immediately after collection, the field soil was sieved to 5 mm to remove debris and then a half of it was immediately stored at 4_J until use as a live soil inoculum. The other half was sterilized with a dose of 25 kGy of 60CO_γ_ irradiation for three days and used as a sterilized soil inoculum. We homogenized 5 g of a slow-release fertilizer (Osmocote Exact Standard, Everris International B.V., Geldermalsen, The Netherlands) with the substrate in each pot. We used gamma irradiation to sterilize the soil as it has been shown to have a limited effect on agronomic properties of air-dry soil (Staunton *et al*. 2002; Wehr & Kirchhof 2021). We separately homogenized 125 ml of the 10 different types of microplastics (five biodegradable and five non-biodegradable) with substrate in each pot, which corresponded to a microplastic concentration of 5% (v/v) in each pot (pot volume was 2.5 L) (**Figure 1**). We applied microplastic concentration of 5% (v/v) as it falls within the range of previously reported microplastic pollution in some soils. For instance, some soils have been shown to contain up to 7% (v/v) of microplastic fragments (Fuller & Gautam., 2016). Moreover, soils in southwestern China were found to contain more than 40,000 microplastic particles per kg of soil (Zhang *et al*. 2018). As control for microplastics, one set of pots did not receive any microplastic treatment (i.e., no-microplastic treatment) and instead received 125 ml of a 1:1 mixture of sand and fine vermiculite.

After transplant, all pots were randomly distributed on four greenhouse benches and their positions were shifted twice (on 18 September 2020 and 7 October 2020) to eliminate any potential bias of variation in environmental conditions in the greenhouse. Plants were watered regularly throughout the experiment. We placed separate plates underneath each pot to hold flow-through water that was then used to rewater the respective pots and prevent a potential cross-contamination of the pots by soil-borne microbes in the pots with different treatments.

### Measurements and calculations

We terminated the experiment on 25 October 2020 (the experiment lasted 55 days) and then immediately harvested shoot biomass of each plant species within each pot separately. The harvest was done in one day. The shoot biomass was dried to a constant biomass for 72h at 65_J and then weighed. To estimate community productivity under the different combinations of microplastic and soil treatments per pot, we summed up dry shoot biomass of all the species per pot. We quantified responses of the plant community diversity to the different combinations of microplastic and soil treatments using inverse of Simpson’s diversity index (Simpson 1949), which estimates the probability of interspecific-encounter if all species are equally abundant (ESN_PIE_) (Harpole *et al*. 2016). The inverse of Simpson’s diversity index takes both species richness and evenness into account and is less sensitive to sampling error and more sensitive to dominant species than the other alpha diversity indices (Hill 1973; Jost *et al*. 2010; Magurran 2021; Rurangwa *et al*. 2021). We applied the following formular to compute the inverse of Simpson’s diversity index:

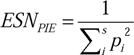

where ESN_PIE_ is an estimate of the inverse of Simpson’s diversity index, *i* = 1, 2… *S*, *P_i_* is the biomass of species *i* in a community divided by the sum of biomass of all species in the community, and *S* is the number of species in the community (Chase & Knight 2013; Harpole e*t al*. 2016; Farrer *et al*. 2016; Liu *et al*. 2019).

We computed ESN_PIE_ values using separate biomass of the different plant species in each experimental pot by using the function *hill_taxa* in the *hillR* package (Jost 2006; Chao, Chiu & Jost 2014). A high value of the inverse of Simpson’s diversity index (ESN_PIE_) denotes high diversity.

### Statistical analyses

To test individual and interactive effects of microplastic types and soil sterilization on plant community productivity indicated by community shoot biomass and community diversity indicated by inverse Simpson’s diversity index, we fitted four Bayesian multilevel models using the function *brm* in the *brms* package (Bürkner 2017) in R 4.0.2 (R Core Team, 2020). In the models, either community productivity or diversity indicators served as dependent variables. To ensure variance normality, we applied a natural-log-transformation to the community productivity data.

We first categorized the 11 microplastic treatment levels into three groups – (1) no- microplastic, (2) biodegradable microplastics, and (3) non-biodegradable microplastics – and examined whether microplastic presence or biodegradability interacted with soil sterilization (*S*) to affect community productivity and diversity. To facilitate this, we created two dummy variables *P1* (no-microplastic *vs.* microplastic [biodegradable and non-biodegradable microplastics combined]) and *P2* (biodegradable microplastics *vs.* non-biodegradable microplastics) to simplify comparisons between groups (sensu Schielzeth 2010). In the two models, the dummy variables *P1* and *P2*, soil treatment (live soil *vs.* sterilized soil), and two-way interaction between soil treatment and *P1* or *P2* were specified as fixed-effect independent variables. To account for non-independence of microplastic types belonging to the same group (i.e., biodegradable or non-biodegradable microplastic) and for non-independence of replicates of the same plant communities, we included individual identity of the microplastics and plant communities as random factors in both models. To relax the homogeneity of variance assumption in the models, we allowed the residual standard deviation *sigma* to vary by the identity of plant communities.

We also fitted two separate models to assess the main and interactive effects of the 10 individual microplastic types and soil treatments on plant community productivity and diversity. In both models, microplastic type, soil treatment, and interaction between them were specified as fixed-effect independent variables. We included identity of the plant communities as a random factor and allowed the residual standard deviation *sigma* to vary by the identity of plant communities for both models. Then, we conducted *post-hoc* comparisons of treatment combinations via the *hypothesis* function in the *brms* package.

In all the four models, we used the default priors set by the *brms* package and ran four independent chains with the *No*- *U*- *Turn Sampler* for 8,000 iterations each (Muth, Oravecz & Gabry 2018), and the first 4,000 iterations were used for warm up. We considered the fixed-effect independent variables as statistically significant when their 95% credible interval of the posterior distribution did not with overlap zero, and as marginally significant when the 90% credible interval did not overlap zero.

## Results

### Plant community biomass

Shoot biomass of the plant communities was significantly influenced by interactive effects of soil treatment and biodegradability of microplastics under averaged across all treatments (significant *S* × *P2*; **Table 1**, **Figure 2A**). Specifically, in the presence of biodegradable microplastics, the mean predict total shoot biomass in living and sterilized soil was 0.60g and 0.26g, respectively. In the presence of non-biodegradable microplastics, the mean predict total shoot biomass in living and sterilized soil was 1.27g and 1.18g, respectively (**Figure 2A**). The model predicted values to find that relative to the effects of non-biodegradable microplastics, biodegradable microplastics had stronger suppressive effects on plant community productivity in sterilized soil (-77.98%) than in live soil (-52.64%) (significant *S*×*P2*; **Figure 2A**). Although soil treatment also had a significant main effect on predict total shoot biomass, but the effect of soil treatment did not significantly interact with the effect of presence of microplastics to influence predict total shoot biomass (non-significant *S* × *P1* interaction; **Table 1 & Figure 2A**).

**Figure 2.**
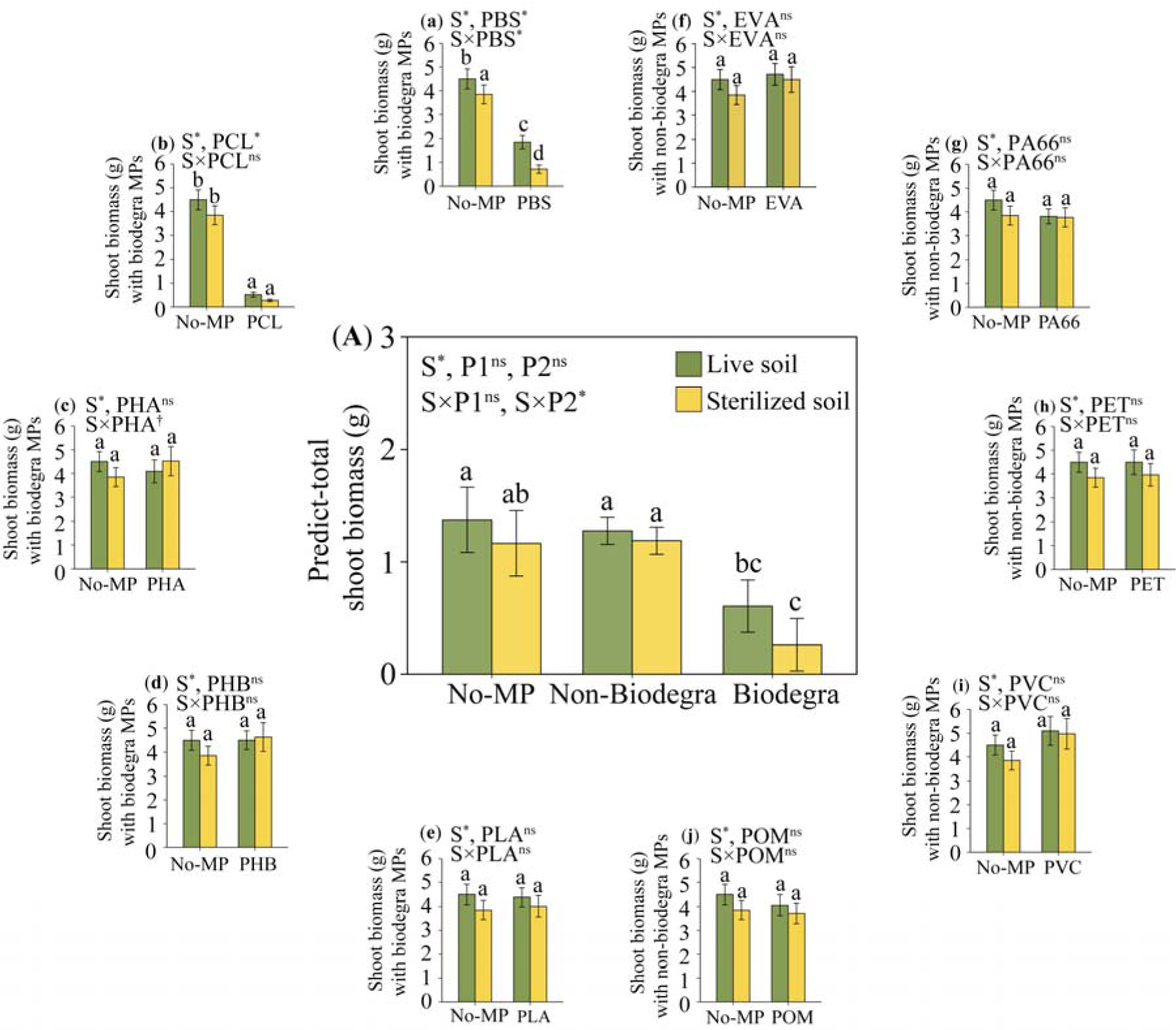
Model-derived mean (±1 SE) estimates of six native plant communities that were grown in the presence of five biodegradable (Biodegra) and five non-biodegradable (Non-biodegra) microplastic types *vs.* absence of microplastics (No-MP) in sterilized soil *vs.* live soil (S). The central panel (A) shows means of communities that were averaged across the 10 microplastics while panels a-j show means that were computed for the individual microplastic types (a-e are for biodegradable and f-j for non-biodegradable microplastics). Asterisks (*) denote statistically significant effects (the 95% credible intervals that do not overlap with zero), while daggers (†) denote marginally significant effects of the treatments (90% credible intervals that do not overlap with zero). Different lowercase letters indicate significant differences among treatments by *post-hoc* mean comparisons.

**Table 1.** The output of Bayesian multilevel models that were run to test the main and interactive effects of soil treatment (sterilized *vs.* live) (*S*), presence of microplastics (microplastics *vs.* no-microplastic) (*P1*), and type of microplastics (biodegradable *vs.* non-biodegradable microplastics) (*P2*) on productivity (indicated by shoot biomass) and diversity (indicated by inverse of Simpson’s diversity index; InvSimpson) of six plant communities. Parameter estimates are statistically significant (marked with asterisks *) if their lower (L) and upper (U) 95% credible intervals (CI) do not overlap with zero and marginally significant (marked with †) if their 90% credible intervals do not overlap with zero.

| Shoot biomass | Estimate | SE | LCI (95%) | UCI (95%) | LCI (90%) | UCI (90%) |
| --- | --- | --- | --- | --- | --- | --- |
| Intercept | **1.120**† | 0.572 | -0.026 | 2.261 | 0.188 | 2.049 |
| *S* | **-0.214** * | 0.054 | -0.318 | -0.109 | -0.303 | -0.124 |
| *P1* | 0.436 | 1.100 | -1.764 | 2.638 | -1.307 | 2.199 |
| *P2* | -0.653 | 0.675 | -2.002 | 0.708 | -1.736 | 0.448 |
| *S* × *P1* | 0.007 | 0.142 | -0.271 | 0.285 | -0.229 | 0.240 |
| *S* × *P2* | **-0.255** * | 0.086 | -0.424 | -0.087 | -0.397 | -0.115 |
| **InvSimpson** |  |  |  |  |  |  |
| Intercept | **1.086** * | 0.086 | 0.91 | 1.258 | 0.952 | 1.224 |
| *S* | **-0.003** * | 0.001 | -0.004 | -0.002 | -0.004 | -0.002 |
| *P1* | 0.004 | 0.006 | -0.007 | 0.015 | -0.005 | 0.013 |
| *P2* | -0.002 | 0.003 | -0.009 | 0.005 | -0.008 | 0.003 |
| *S* × *P1* | **-0.004** * | 0.002 | -0.007 | -0.001 | -0.006 | -0.001 |
| *S* × *P2* | -0.001 | 0.001 | -0.002 | 0.001 | -0.002 | 0.001 |

Residual standard deviation sigma for the six separate communities are presented in Table S2.

In the separate analyses of the main and interactive effects of soil and the 10 individual microplastics, the community shoot biomass was only significantly influenced by an interaction between the biodegradable microplastic PBS and soil treatment (**Figure 2a & Table S3**). Specifically, relative to the shoot biomass of communities that were grown in the absence of microplastic (No-MP), the presence of PBS caused a greater decline in mean community shoot biomass in sterilized soil (-81.42%) than in live soil (-58.79%) (**Figure 2a**). Although we found that compared to mean shoot biomass of communities that were grown in No-MP, presence of PHA slightly decreased mean community shoot biomass in live soil (-9.04%) but tended to increase (+17.53%) biomass in sterilized soil, the interactive effect between PHA and soil treatment was not statistically significant (**Figure 2c & Table S3**).

### Plant community diversity

Soil treatment significantly interacted with the presence of microplastics to influence native plant community diversity (significant *S × P1* ; Table 1 & Figure 3A). Relative to the diversity of native plant communities that were grown in No-MP, presence of either biodegradable or non-biodegradable microplastics caused a significant reduction in the diversity of native plant communities in live soil (-23.68%), but there was no significant effect on plant diversity in sterilized soil (-1.84%) (significant S×P1 ; Table 1 & Figure 3A). However, either biodegradable or non-biodegradable microplastics both had negligible effects on the native plant community in the sterilized soil (non- significant *S*× *P2* ; **Figure 3A & Table 1**).

**Figure 3.**
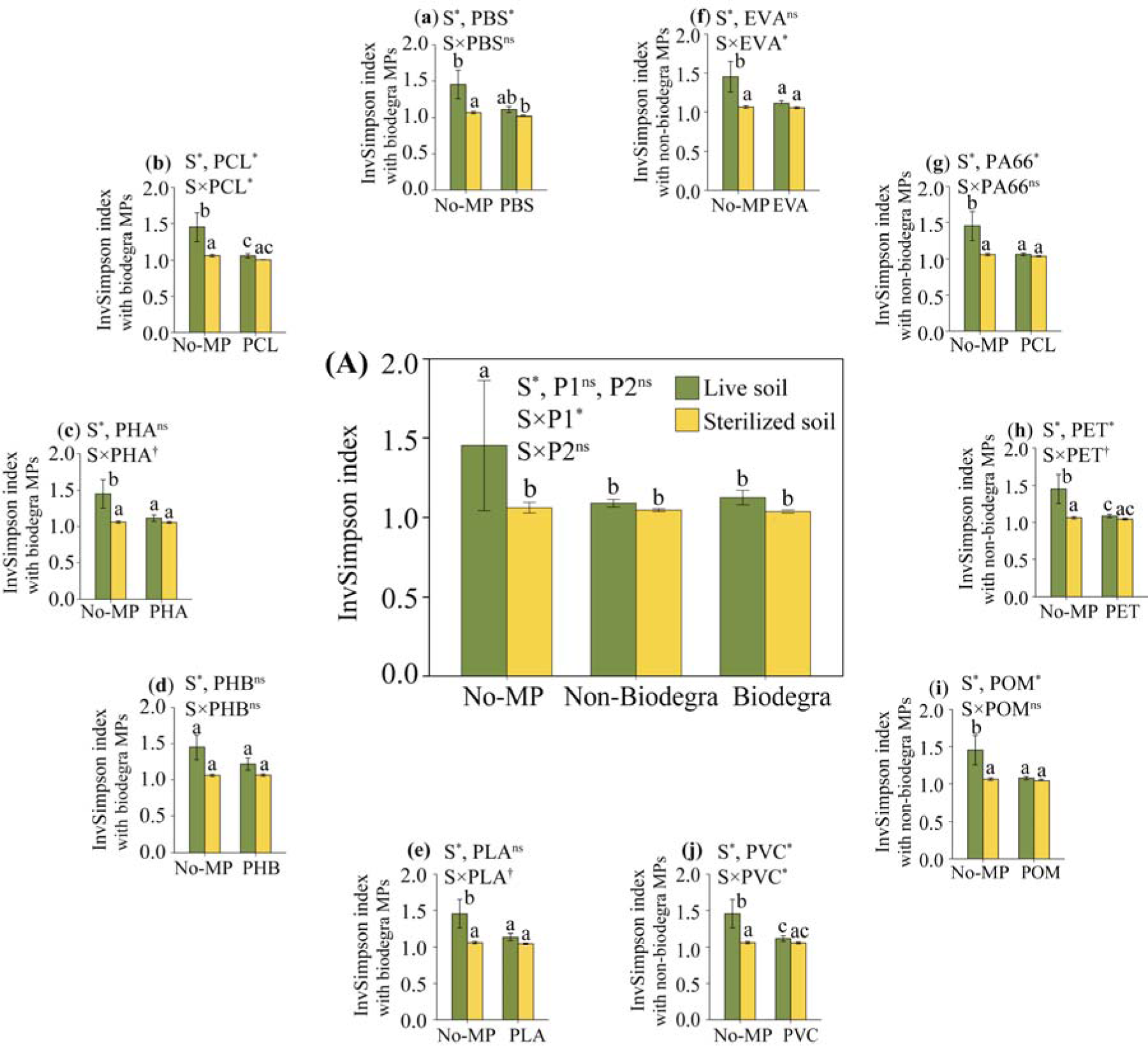
Mean (± 1SE) of diversity (indicated by inverse of Simpson’s diversity index; InvSimpson) of six native plant communities indicated by inverse Simpson index (InvSimpson) that were grown in the presence of five biodegradable (Biodegra) and five non-biodegradable (Non-biodegra) microplastic types *vs.* absence of microplastics (No-MP) in sterilized soil *vs.* live soil (S). The central panel (A) shows means of communities that were averaged across the 10 microplastics while panels a-j show means that were computed for the individual microplastic types (a-e are for biodegradable and f-j for non-biodegradable microplastics). Asterisks (*) denote statistically significant effects (the 95% credible intervals that do not overlap with zero), while daggers (†) denote marginally significant effects of the treatments (90% credible intervals that do not overlap with zero). Different lowercase letters indicate significant differences among treatments by *post-hoc* mean comparisons.

In the separate analyses of the main and interactive effects of the 10 individual microplastics and soil treatment, the community diversity was significantly influenced by an interaction between the PCL, EVA, PVC and soil treatment. Specifically, relative to the community diversity that were grown in No-MP, the presence of PCL caused a greater decline in mean community diversity in living soil (-27.03%) than in sterilized soil (-5.28%) (**Figure 3b**), and the trends for EVA (living soil, -23.46%; sterilized soil, -0.83%) and PVC (living soil, -23.06%; sterilized soil, -0.31%) were as for PCL (**Figure 3f, j)**. There were marginally significant interactions between soil treatment and the biodegradable microplastics PHA and PLA (S×PHA and S×PLA) (**Figure 3c, e & Table S4**) and non-biodegradable microplastics PET (S×PET) on community diversity (**Figure 3h & Table S4**).

## Discussion

Averaged across all treatments, the main effects of microplastics addition (absence or presence), and microplastics type (biodegradable *vs.* non-biodegradable) on plant community productivity (indicated by community shoot biomass) and diversity (indicated by inverse of Simpson’s diversity index) were both not significant (**Table 1**, **Figures 2A & Figures 3A**). Instead, plant community productivity and diversity were significantly influenced by interactive effects of soil and microplastic treatments. Notably, the model predicted values to find that relative to the effects of non-biodegradable microplastics, biodegradable microplastics had stronger suppressive effects on plant community productivity in sterilized soil (-77.98%) than in live soil (-52.64%) (significant *S*×*P2*; **Figure S1**). This finding suggests that soil biota in the live soil ameliorated suppressive effects of biodegradable microplastics on plant community productivity under averaged across all treatments. Furthermore, by analysing the effect of each microplastic on total shoot biomass, we found that the individual microplastic PBS largely drove the overall pattern noted here. In contrast, the presence of microplastics (regardless of biodegradable or non-biodegradable microplastics) suppressed plant community diversity more strongly in live soil than in sterilized soil under averaged across all treatments (significant *S*×*P1*, **Figure 3A**). Overall, these results suggest that soil biota may weaken negative effects of biodegradable microplastics on native plant community productivity, while the opposite was observed for soil biota modulation of plant community diversity in the presence of microplastics under averaged across all treatments.

Among the 10 individual microplastics, the biodegradable microplastic PBS had a significantly less negative impact on the total community shoot biomass in live soil than sterilized soil (**Figure 2a**), which indicates that soil biota in the live soil partly degraded the PBS and made it less harmful to the plants. Studies have shown that soil fungi and bacteria can degrade microplastics into less harmful substances and that different kinds of microplastics differ in the rate of decay (Yuan *et al*. 2020). The microbes use microplastics as a carbon source (Yuan *et al*. 2020). A previous study found that strain WF-6 of the fungus *Fusarium solani* that was isolated from a farmland degraded 2.8% of the PBS over a period of 14 days (Abe *et al*. 2010). Other species of fungi (*Aspergillus versicolor, A. fumigatus,* and *Penicillium*) and bacteria (*Bacillus* and *Thermopolyspora*) have also been shown to degrade PBS (Zhao *et al*. 2005; Ishii *et al*. 2008). Biodegradation rates of the other microplastics that we studied presently have also been studied. For instance, the fungus *Pullularia pullulans* degraded PCL at the rate of 16 mg/cm^2^ surface area at 30°C (Fields *et al*. 1974). Another study compared biodegradation of PCL, PHB, PLA, and PBS in soil and compost over a period of more than 10_Jmonths at 25_J°C, 37 °C and 50_J°C (Hosni *et al*. 2019). The study found that biodegradation rates varied among the microplastic types and incubation temperatures, although PCL showed the fastest biodegradation rate under all conditions and was completely degraded when buried in compost and incubated at 50 °C after 91_Jdays (Hosni *et al*. 2019). Although we did not compare biodegradation rates of the 10 microplastics, it is likely that PBS had the highest rate of decay in the live soil that we used in the present study. Future studies may compare degradation rates of the 10 microplastics in the soil that we used in the present experiment and the specific soil microorganisms involved in the biodegradation.

The present finding that PBS significantly suppressed plant community productivity (**Figure 2a**) contrasts with that of another study that found that PBS did not have adverse effects on the growth of *Brassica rapa* (Chinese cabbage) (Wang *et al*. 2019). However, in agreement with the present finding where the biodegradable microplastic PHB did not significantly influence total shoot biomass of plant communities (**Figure 2d**), a previous study did not find significant adverse effect of PHB on growth of *Zea mays* (maize) (Kontárová *et al*. 2022). In contrast to the present findings of non-significant effects of non-biodegradable microplastics PET and PVC on the plant communities (**Figure 2h & i**), other studies found inhibitory effects of PET on *Lepidium sativum* (garden cress) (Pignattelli *et al*. 2021) and PVC on *Triticum aestivum* (wheat) (Xinyue *et al*. 2021). In general, these results suggest that the impact of microplastics on plant growth may depend upon a range of factors including plant growth environment, plant species identity, as well as whether a plant grows in a community context or not.

The presence of microplastics (regardless of biodegradable or non-biodegradable microplastics) suppressed plant community diversity more strongly in live soil than in sterilized soil under averaged across all treatments (**Figure 3A)**. Furthermore, we found that this pattern held for the microplastics PCL, EVA, and PVC (**Figure 3b, h & j**), which suggests that these individual microplastics idendities largely drove the overall pattern here. It is likely that microorganisms present in the live soil mediated the impacts of the microplastics on plant community diversity. In particular, the microplastics may have suppressed diversity, abundance, and activities of soil-borne microbial mutualists of some plant species in the plant communities, which then fed back to negatively affect diversity of the plant communities. Additionally, or alternatively, the microplastics may have stimulated growth and activities of some soil-borne microbial antagonists of the plant species, which then suppressed diversity of the plant communities. Indeed, empirical studies have shown that microplastics, depending on their properties such as biodegradability, hydrophobicity, electric charge, or roughness, can alter community composition and enzymatic activities of beneficial and pathogenic microorganisms in the soil (Wang *et al*. 2016). In one study, an experimental addition of polyethylene microplastics into the soil remarkably altered bacterial community structure, induced accumulation of pathogens, and caused an increase in the urease and catalase activities in the soil (Huang *et al*. 2019). Separately, addition of polyethylene and PVC microplastics into the soil influenced enzymatic activities, abundance and diversity in bacteria (Fei *et al*. 2020). Specifically, the microplastics inhibited activities of fluorescein diacetate hydrolase, stimulated the activities of urease and acid phosphatase, increased abundance of *Burkholderiaceae* but decreased richness and diversity of *Sphingomonadaceae* and *Xanthobacteraceae* bacteria (Fei *et al*. 2020). Microplastics can also alter diversity, abundance, and activities of plant pathogenic and beneficial soil fungi (Yu *et al*. 2021; Fan, Tan & Yu 2022), microarthropod and nematode communities (Lin *et al*. 2020). By altering the community composition and activities of soil microorganisms, microplastics may affect plant growth indirectly through alteration in soil carbon and nutrient cycling, altered soil water and nutrient acquisition by plants, and accumulation of pathogens on plants. Future studies may unravel which of these mechanisms were at play in our study system.

Results of the present study complement those of other studies that tested the effects of microplastics on plant growth performance, with variable outcomes. For instance, the present results partly support those of Lozano & Rillig (2020) who noted that addition of a non-biodegradable microplastic in the form of polyester fibers into the soil enhanced biomass production by *Calamagrostis epigejos* but simultaneously suppressed biomass of *Holcus lanatus* when the plants were grown in a plant community consisting of seven plant species. In other studies, *Lolium perenne* (perennial ryegrass) showed reduced biomass after exposure to the biodegradable microplastic PLA (Boots, Russell & Green 2019). In contrast, exposure to the non-biodegradable microplastics polyester and polystyrene increased *Allium fistulosum* (spring onion) root biomass, while polyethylene high density, PET, and polypropylene microplastics had only weak effects on the plant (de Souza Machado *et al*. 2019). As the previous studies did not investigate whether soil microbial communities can influence the effects of microplastics on plant community growth and diversity, much uncertainty remains regarding the ability to predict soil microbe-mediated effects of microplastics on plant community productivity and structure in natural habitats.

Although the specific taxa of microorganisms that likely drove the process were not identified, several studies have identified different groups of fungi and bacteria that produce various intracellular and extracellular enzymes with the ability to break down biodegradable microplastics, and biodegradability of the microplastics ranged from 13% to 100% over a period of 18-300 days (Emadian, Onay & Demirel 2017). Variability in biodegradability of microplastics has been ascribed to various factors including variation in the physical and chemical structure of the microplastics, the type of enzyme produced by microorganisms, and environmental conditions like pH, temperature, moisture, and oxygen content of the medium in which biodegradation takes place (Boyandin *et al*. 2013; Emadian, Onay & Demirel 2017). For instance, two types of PHAs, the homopolymer 3-hydroxybutyric acid and the copolymer of 3-hydroxybutyric/3-hydroxyvaleric acids, had variable rates of biodegradation in soil at two Vietnamese locations with contrasting soil and climatic conditions (Boyandin *et al*. 2013). In particular, the homopolymer degraded at higher rates than the copolymer at one location (Boyandin *et al*. 2013). The two study locations also harbored markedly different soil bacterial and fungal communities that were linked to the different rates of biodegradation of the PHAs (Boyandin *et al*. 2013). Future studies may identify the specific taxa of soil microorganisms that may have degraded the microplastics that we studied, their rates of biodegradation, and the effects thereof on plant community structure and productivity under more natural field conditions in contrasting climatic conditions.

## Conclusions

Our multi-species experiments revealed that the addition of microplastics had different effects on plant community productivity and diversity in live *vs.* sterilized soil. Particularly, live soil ameliorated the negative effects of biodegradable microplastics on total shoot biomass of the plant communities under averaged across all treatments, and further analyses revealed that the individual microplastic PBS largely drove the overall pattern noted here. In contrast, for the plant community diversity, presence rather than type (biodegradable *vs.* non-biodegradable) of microplastics influenced community diversity suppressed plant community diversity more strongly in live soil than in sterilized soil, and this tendency was also strongly conditioned by few polymer microplastics. Overall, our findings indicate that even biodegradable microplastics, which are considered environmentally friendly, still pose significant ecological risks to the structure and productivity of plant communities with potential implications for functioning of terrestrial ecosystems. More studies on the effects of biodegradable and non-biodegradable microplastics on plant community productivity and diversity under a wide range of biotic and abiotic conditions may reveal the general pattern of impacts of microplastics on plant communities and ecosystem processes.

## Acknowledgement

We thank Lingxi Wang, Xue Zhang, Yimin Yan, Zhengkuan Lu and Lichao Wang for assistance with the experiment. Yanjie Liu acknowledges funding from the Chinese Academy of Sciences (Y9B7041001). Yanmei Fu acknowledges funding from the National Natural Science Foundation of China (32201341). Ayub M.O. Oduor was supported by a fellowship of the Chinese Academy of Sciences - President’s International Fellowship Initiative (CAS-PIFI) (2021VBB0004).

## Author contributions

**Yanjie Liu** conceived the idea. **Ming Jiang and Yanjie** Liu designed the experiment. **Yanmei Fu** performed the experiment. **Yanmei Fu, Ayub. M. O. Oduor** and **Yanjie Liu** analyzed the data. **Yanmei Fu** and **Ayub. M. O. Oduor** wrote the draft of the manuscript, with further input from **Ming Jiang** and **Yanjie Liu**.

## Data accessibility

Should the manuscript be accepted, the data supporting the results will be archived in Dryad and the data DOI will be included at the end of the article.

## Data table

