## Supplemental Tables for "Live soil ameliorated the negative effects of biodegradable but not non-biodegradable microplastics on the growth of plant communities"

**Supporting information**

**Table S1.** Details of 28 plant species that were used in the current experiment.

| **Species** | **Family** | **Community group** | **Sowing date** | **Life cycle** | **Seed source** |
| --- | --- | --- | --- | --- | --- |
| *Rumex crispus* L. | Polygonacea | Native Community 1 | 07.08.2020 | Perennials | Nei Mongol, China |
| *Suaeda prostrata* Pall. | Amaranthaceae | Native Community 1 | 07.08.2020 | Annual | Nei Mongol, China |
| *Artemisia rubripes* Nakai | Compositae | Native Community 1 | 11.08.2020 | Perennials | Nei Mongol, China |
| *Elymus sibiricus* Linn. | Poaceae | Native Community 1 | 14.08.2020 | Perennials | Nei Mongol, China |
| *Hordeum brevisubulatum* (Trin.) Link | Poaceae | Native Community 1 | 14.08.2020 | Perennials | JiLin, China |
| *Arabis pendula* L. | Cruciferae | Native Community 2 | 07.08.2020 | Biennial | Nei Mongol, China |
| *Artemisia stechmanniana* Bess. | Compositae | Native Community 2 | 11.08.2020 | Perennials | Nei Mongol, China |
| *Cynoglossum divaricatum* Stephan ex Lehm. | Boraginaceae | Native Community 2 | 11.08.2020 | Perennials | Nei Mongol, China |
| *Setaria arenaria* Kitag. | Poaceae | Native Community 2 | 14.08.2020 | Annual | Nei Mongol, China |
| *Koeleria macrantha* | Poaceae | Native Community 2 | 07.08.2020 | Perennials | Nei Mongol, China |
| *Crepidiastrum sonchifolium* (Maximowicz) Pak & Kawano | Compositae | Native Community 3 | 07.08.2020 | Annual/Biennial | Nei Mongol, China |
| *Potentilla tanacetifolia* Willd. ex Schlecht. | Rosaceae | Native Community 3 | 07.08.2020 | Perennials | Nei Mongol, China |
| *Geum aleppicum* Jacq. | Rosaceae | Native Community 3 / 6 | 07.08.2020 | Perennials | Nei Mongol, China |
| *Digitaria sanguinalis* (L.) Scop. | Poaceae | Native Community 3 | 07.08.2020 | Annuals | JiLin, China |
| *Echinochloa crus-galli* | Poaceae | Native Community 3 | 07.08.2020 | Annuals | JiLin, China |
| *Potentilla longifolia* Willd. ex Schlecht. | Rosaceae | Native Community 4 | 07.08.2020 | Perennials | Nei Mongol, China |
| *Gypsophila licentiana* Hand.-Mazz. | Caryophyllaceae | Native Community 4 | 07.08.2020 | Perennials | Nei Mongol, China |
| *Dontostemon micranthus* C. A. Mey. | Brassicaceae | Native Community 4 | 07.08.2020 | Annual/Biennial | Nei Mongol, China |
| *Setaria pumila* (Poir.) Roem. & Schult | Poaceae | Native Community 4 / 6 | 14.08.2020 | Annuals | Nei Mongol, China |
| *Chloris virgata* Sw*.* | Poaceae | Native Community 4 | 14.08.2020 | Annuals | Nei Mongol, China |
| *Malva verticillata* var. *crispa* | Malvaceae | Native Community 5 | 14.08.2020 | Annuals | JiLin, China |
| *Cynanchum chinense* R.Br. | Apocynaceae | Native Community 5 | 14.08.2020 | Annuals | Nei Mongol, China |
| *Euphorbia humifusa* Willd | Euphorbiaceae | Native Community 5 | 14.08.2020 | Annuals | JiLin, China |
| *Agrostis matsumurae* Hack. ex Honda | Poaceae | Native Community 5 | 11.08.2020 | Perennials | Nei Mongol, China |
| *Leymus secalinus* (Georgi)Tzvel. | Poaceae | Native Community 5 | 11.08.2020 | Perennials | Nei Mongol, China |
| *Achillea asiatica* Serg. | Compositae | Native Community 6 | 07.08.2020 | Annuals | Nei Mongol, China |
| *Poa annua* L. | Poaceae | Native Community 6 | 07.08.2020 | Annuals | Nei Mongol, China |
| *Cleistogenes squarrosa* (Trin.) Keng | Poaceae | Native Community 6 | 11.08.2020 | Perennials | JiLin, China |

**Table S2.** Output of the Bayesian multilevel model estimates of the residual standard-deviation sigma for each plant community in Table1. Shown are the model estimates,standard errors (SE) as well as 95% and 90% lower (L) and upper (U) credible intervals (CI).

|  | **Community group** | **Estimate** | **SE** | **LCI (95%)** | **UCI (95%)** | **LCI (90%)** | **UCI (90%)** |
| --- | --- | --- | --- | --- | --- | --- | --- |
| **Shoot biomass** | sigma_groupA | **-0.821*** | 0.082 | -0.977 | -0.655 | -0.952 | -0.683 |
|  | sigma_groupB | **-1.020*** | 0.085 | -1.182 | -0.848 | -1.156 | -0.877 |
|  | sigma_groupC | **-0.575*** | 0.079 | -0.724 | -0.415 | -0.702 | -0.443 |
|  | sigma_groupD | **-0.644*** | 0.081 | -0.796 | -0.485 | -0.774 | -0.510 |
|  | sigma_groupE | **-0.773*** | 0.082 | -0.930 | -0.605 | -0.904 | -0.635 |
|  | sigma_groupF | **-1.012*** | 0.084 | -1.170 | -0.842 | -1.147 | -0.869 |
| **InvSimpson** | sigma_groupA | **-0.617*** | 0.077 | -0.763 | -0.459 | -0.739 | -0.487 |
|  | sigma_groupB | **-3.682*** | 0.077 | -3.827 | -3.525 | -3.805 | -3.553 |
|  | sigma_groupC | **-4.213*** | 0.078 | -4.360 | -4.056 | -4.338 | -4.081 |
|  | sigma_groupD | **-5.767*** | 0.113 | -5.985 | -5.543 | -5.950 | -5.580 |
|  | sigma_groupE | **-4.338*** | 0.078 | -4.486 | -4.178 | -4.463 | -4.208 |
|  | sigma_groupF | **-5.874*** | 0.097 | -6.057 | -5.676 | -6.029 | -5.710 |

**Table S3.** The output of Bayesian multilevel models that were run to test the main and interactive effects of soil treatment (sterilized soil *vs.* live soil) (*S*) and 10 individual microplastic types on productivity (indicated by total shoot biomass) of six plant communities. Separate models were run for each microplastic. Parameter estimates are statistically significant (marked with asterisks *) if their lower (L) and upper (U) 95% credible intervals (CI) do not overlap with zero, and marginally significant (marked with †) if their 90% credible intervals do not overlap with zero.

|  | **Estimate** | **SE** | **LCI (95%)** | **UCI (95%)** | **LCI (90%)** | **UCI (90%)** |
| --- | --- | --- | --- | --- | --- | --- |
| *Intercept* | **0.889*** | 0.403 | 0.084 | 1.702 | 0.249 | 1.529 |
| *S* | **-0.217*** | 0.039 | -0.293 | -0.142 | -0.281 | -0.154 |
| *PBS* | **-1.593*** | 0.093 | -1.775 | -1.410 | -1.744 | -1.441 |
| *PCL* | **-2.586*** | 0.100 | -2.778 | -2.387 | -2.748 | -2.421 |
| *PHA* | -0.039 | 0.088 | -0.210 | 0.138 | -0.183 | 0.109 |
| *PHB* | 0.046 | 0.089 | -0.126 | 0.223 | -0.098 | 0.191 |
| *PLA* | 0.015 | 0.088 | -0.158 | 0.189 | -0.131 | 0.161 |
| *EVA* | 0.055 | 0.090 | -0.119 | 0.229 | -0.091 | 0.203 |
| *PA66* | -0.080 | 0.089 | -0.254 | 0.095 | -0.225 | 0.066 |
| *PET* | -0.037 | 0.088 | -0.209 | 0.138 | -0.182 | 0.108 |
| *POM* | -0.149 | 0.091 | -0.326 | 0.032 | -0.296 | 0.003 |
| *PVC* | 0.024 | 0.089 | -0.151 | 0.199 | -0.124 | 0.169 |
| *S×PBS* | **-1.066*** | 0.181 | -1.420 | -0.712 | -1.359 | -0.770 |
| *S×PCL* | -0.211 | 0.177 | -0.560 | 0.136 | -0.502 | 0.080 |
| *S×PHA* | **0.294†** | 0.176 | -0.051 | 0.637 | 0.005 | 0.581 |
| *S×PHB* | 0.116 | 0.177 | -0.230 | 0.461 | -0.173 | 0.404 |
| *S×PLA* | 0.083 | 0.176 | -0.258 | 0.431 | -0.206 | 0.375 |
| *S×EVA* | 0.161 | 0.178 | -0.190 | 0.510 | -0.131 | 0.452 |
| *S×PA66* | 0.138 | 0.176 | -0.203 | 0.482 | -0.147 | 0.423 |
| *S×PET* | 0.046 | 0.175 | -0.294 | 0.385 | -0.242 | 0.332 |
| *S×POM* | 0.044 | 0.177 | -0.306 | 0.387 | -0.246 | 0.332 |
| *S×PVC* | 0.137 | 0.181 | -0.220 | 0.489 | -0.161 | 0.436 |

**Table S4.** The output of Bayesian multilevel models that were run to test the main and interactive effects of soil treatment (sterilized soil *vs.* live soil) (*S*) and 10 individual microplastic types on structure uniformity of six plant communities indicated byinverse of Simpson's diversity index (InvSimpson). Parameter estimates are statistically significant (marked with asterisks *) if their lower (L) and upper (U) 95% credible intervals (CI) do not overlap with zero and marginally significant (marked with †) if their 90% credible intervals do not overlap with zero.

|  | **Estimate** | **SE** | **LCI (95%)** | **UCI (95%)** | **LCI (90%)** | **UCI (90%)** |
| --- | --- | --- | --- | --- | --- | --- |
| *Intercept* | **1.0840*** | 0.0908 | 0.9045 | 1.2724 | 0.9482 | 1.2244 |
| *S* | **-0.0020*** | 0.0004 | -0.0032 | -0.0014 | -0.0030 | -0.0016 |
| *PBS* | **-0.0110*** | 0.0012 | -0.0134 | -0.0087 | -0.0130 | -0.0091 |
| *PCL* | **-0.0130*** | 0.0013 | -0.0151 | -0.0102 | -0.0147 | -0.0105 |
| *PHA* | -0.0011 | 0.0010 | -0.0031 | 0.0009 | -0.0027 | 0.0006 |
| *PHB* | -0.0009 | 0.0010 | -0.0029 | 0.0011 | -0.0026 | 0.0008 |
| *PLA* | -0.0007 | 0.0010 | -0.0028 | 0.0013 | -0.0024 | 0.0010 |
| *EVA* | -0.0009 | 0.0011 | -0.0031 | 0.0013 | -0.0027 | 0.0009 |
| *PA66* | **-0.0030*** | 0.0010 | -0.0046 | -0.0006 | -0.0043 | -0.0009 |
| *PET* | **-0.0030*** | 0.0010 | -0.0048 | -0.0008 | -0.0045 | -0.0011 |
| *POM* | **-0.0050*** | 0.0011 | -0.0072 | -0.0030 | -0.0068 | -0.0033 |
| *PVC* | **-0.0020*** | 0.0010 | -0.0043 | -0.0002 | -0.0040 | -0.0006 |
| *S×PBS* | 0.0028 | 0.0021 | -0.0012 | 0.0069 | -0.0006 | 0.0063 |
| *S×PCL* | **0.0050*** | 0.0021 | 0.0008 | 0.0089 | 0.0014 | 0.0082 |
| *S×PHA* | **0.0040†** | 0.0021 | -0.0001 | 0.0080 | 0.0005 | 0.0073 |
| *S×PHB* | 0.0014 | 0.0021 | -0.0026 | 0.0054 | -0.0020 | 0.0047 |
| *S×PLA* | **0.004†** | 0.0021 | -0.0003 | 0.0077 | 0.0003 | 0.0071 |
| *S×EVA* | **0.005*** | 0.0021 | 0.0009 | 0.0090 | 0.0015 | 0.0083 |
| *S×PA66* | 0.0030 | 0.0020 | -0.0010 | 0.0070 | -0.0003 | 0.0063 |
| *S×PET* | **0.004†** | 0.0020 | -0.0004 | 0.0076 | 0.0003 | 0.0069 |
| *S×POM* | 0.0024 | 0.0021 | -0.0016 | 0.0065 | -0.0010 | 0.0058 |
| *S×PVC* | **0.0060*** | 0.0021 | 0.0016 | 0.0096 | 0.0022 | 0.0090 |
